# _CG_-SLENP: A chemical genetics strategy to selectively label existing proteins and newly synthesized proteins

**DOI:** 10.1101/2023.11.02.565346

**Authors:** Jian Wang, Bo Chao, Jake Piesner, Felice Kelly, Stefanie Kaech Petrie, Xiangshu Xiao, Bingbing X. Li

## Abstract

Protein synthesis and subsequent delivery to the target locations in cells are essential for their proper functions. Methods to label and distinguish newly synthesized proteins from existing ones are critical to assess their differential properties, but such methods are lacking. We describe the first <u>c</u>hemical <u>g</u>enetics-based approach for <u>s</u>elective labeling of <u>e</u>xisting and <u>n</u>ewly synthesized <u>p</u>roteins that we termed as _CG_-SLENP. Using HaloTag in-frame fusion with lamin A (LA), we demonstrate that the two pools of proteins can be selectively labeled using _CG_-SLENP in living cells. We further employ our recently developed selective small molecule ligand **LBL1** for LA to probe the potential differences between newly synthesized and existing LA. Our results show that **LBL1** can differentially modulate these two pools of LA. These results indicate that the assembly states of newly synthesized LA are distinct from existing LA in living cells. The _CG_-SLENP method is potentially generalizable to study any cellular proteins.

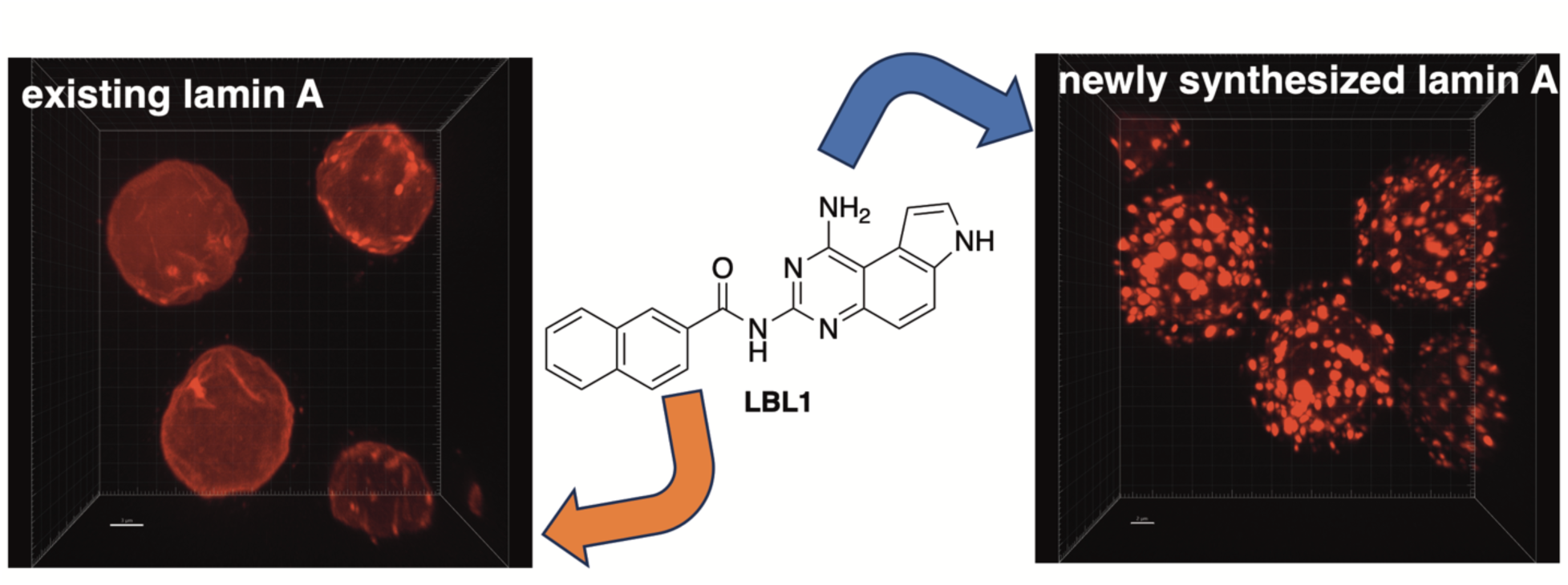

## INTRODUCTION

The cellular protein synthesis is essential for protein homeostasis and life. Once proteins are synthesized at the ribosomes, they undergo different posttranslational modifications and trafficking processes to reach their target destinations to execute their functions in cells. Almost all proteins in cells execute their functions through formation of specialized multimeric complexes.^1^ One mechanism for the formation of multimeric protein complexes is through random or directed collision to form complexes once the protein subunits are fully synthesized and released from the ribosomes. Alternatively and perhaps more likely, the protein subunits are folded and the protein complexes are formed while they are being translated at the ribosomes.^2, 3^ This so-called co-translational assembly mechanism offers potential benefits of stabilizing nascent polypeptide chains and preventing formation of toxic polypeptides, which in turn facilitates the requisite complex formation.^2, 4^ However, the nature and properties of newly formed complexes in cells are largely unknown partly due to the significant challenges associated with identifying, isolating and characterizing these complexes in cells.

Mammalian cells have an intricate network of proteins that form the cytoskeleton. These include microtubules, actin filaments and various intermediate filaments (IF).^5^ Among the IF proteins, nuclear lamins are type V IF proteins localized underneath inner nuclear membrane (INM).^6^ In mammalian cells, three lamin genes lamin A, B1, B2 (*LMNA*, *LMNB1* and *LMNB2*) encode four major lamin proteins: lamin A (LA), lamin B1 (LB1), lamin B2 (LB2) and lamin C (LC) that form the proteinaceous meshwork underneath INM.^7^ All the lamin proteins contain an *N*-terminal non-helical head domain, a central long coiled-coil domain, a nuclear localization sequence (NLS) and a *C*-terminal globular immunoglobulin (Ig) fold.^8^ Through the central coiled-coil domain, lamins form dimers that are further assembled into long filaments with the originally estimated diameter of ∼10 nm.^9^ More recent cryo-electron tomography (cryo-ET) studies indicated that the diameter of lamins in somatic cells is ∼3.5 nm, which is significantly smaller than initially estimated.^10, 11^

As a nuclear protein, LA is synthesized in the cytosol and then translocated into nucleus to be facilitated by the NLS in LA using nuclear importin receptors.^12^ Many nuclear proteins including nuclear transcription complexes and nuclear pore complexes have been reported to undergo co-translational complex formation for subsequent nuclear translocation.^13, 14^ It has also been suggested that LA is dimerized in the cytosol either as a homodimer or heterodimer for subsequent nuclear translocation.^12^ However, the nature about the higher-order structures of newly synthesized LA relative to existing LA in the nuclear envelope is unknown. In this paper, we designed the first _CG_-SLENP strategy to distinguish the newly synthesized LA from existing LA. We further employed small molecule ligand **LBL1** to evaluate its effect on the dynamics of newly synthesized and existing LA pools using _CG_-SLENP, providing fresh insights into the assembly states of LA in cells. **LBL1** is the first small molecule ligand known to selectively bind to nuclear lamins and it can stabilize oligomeric state of LA.^15, 16^

## RESULTS AND DISCUSSION

In order to investigate the nature of newly synthesized proteins, we would need a SLENP method to distinguish this population from the existing counterpart. Previously described metabolic labeling methods using radioactive amino acids or bioorthogonally modified amino acids^17, 18^ can lead to identification of newly synthesized species, but not the existing one. This metabolic labeling approach also involves laborious downstream procedures including immunoprecipitation to identify the specific protein of interest. To circumvent these issues, we developed a chemical genetics strategy with the goal to selectively label newly synthesized and existing proteins. By taking LA as an example, our _CG_-SLENP strategy is depicted in Figures 1 and 2. We envisioned that expression of HaloTag-LA fusion in cells would help distinguish the two populations using different chloroalkane ligands. HaloTag is an engineered bacterial dehalogenase that covalently reacts with chloroalkane compounds, which are collectively called halo ligands.^19^ We hypothesized that upon reacting with a silent halo ligand to exhaust existing HaloTag-LA fusion, the newly synthesized HaloTag-LA can be labeled by an orthogonal halo ligand to effectively distinguish the two populations (Figure 2A). Thus, a lentiviral construct expressing HaloTag protein in-frame with LA fusion was created and sequence verified (see Figure S1 for sequence). This construct was then stably expressed in MDA-MB-468 cells using lentiviruses. As shown in Figure 1B, the fusion was expressed in cells at a similar level to the endogenous LA protein. As reported before by us and others in different cell types,^15, 20^ endogenous LA was stably integrated into nuclear lamina underneath INM in MDA-MB-468 cells (Figure 1C, top). At the same time, a fraction of LA was also detected in the nucleoplasm. To examine if exogenously expressed HaloTag-LA was also integrated into the nuclear lamina, we treated MDA-MB-468 cells expressing HaloTag-LA with a fluorescent Janelia Fluor Halo ligand (Halo-JF549) (Figure 1A). The cells were also co-stained with anti-LA. As shown in Figure 1C (bottom), the Halo ligand signal was very similar to the LA signal in cells. We did, however, observe more nucleoplasmic LA signal in these cells, compared to parental cells. Treating the parental cells with Halo-JF549 did not generate any fluorescent signal (Figure 1C, top), supporting the specificity of the Halo ligand.

**Figure 1.**
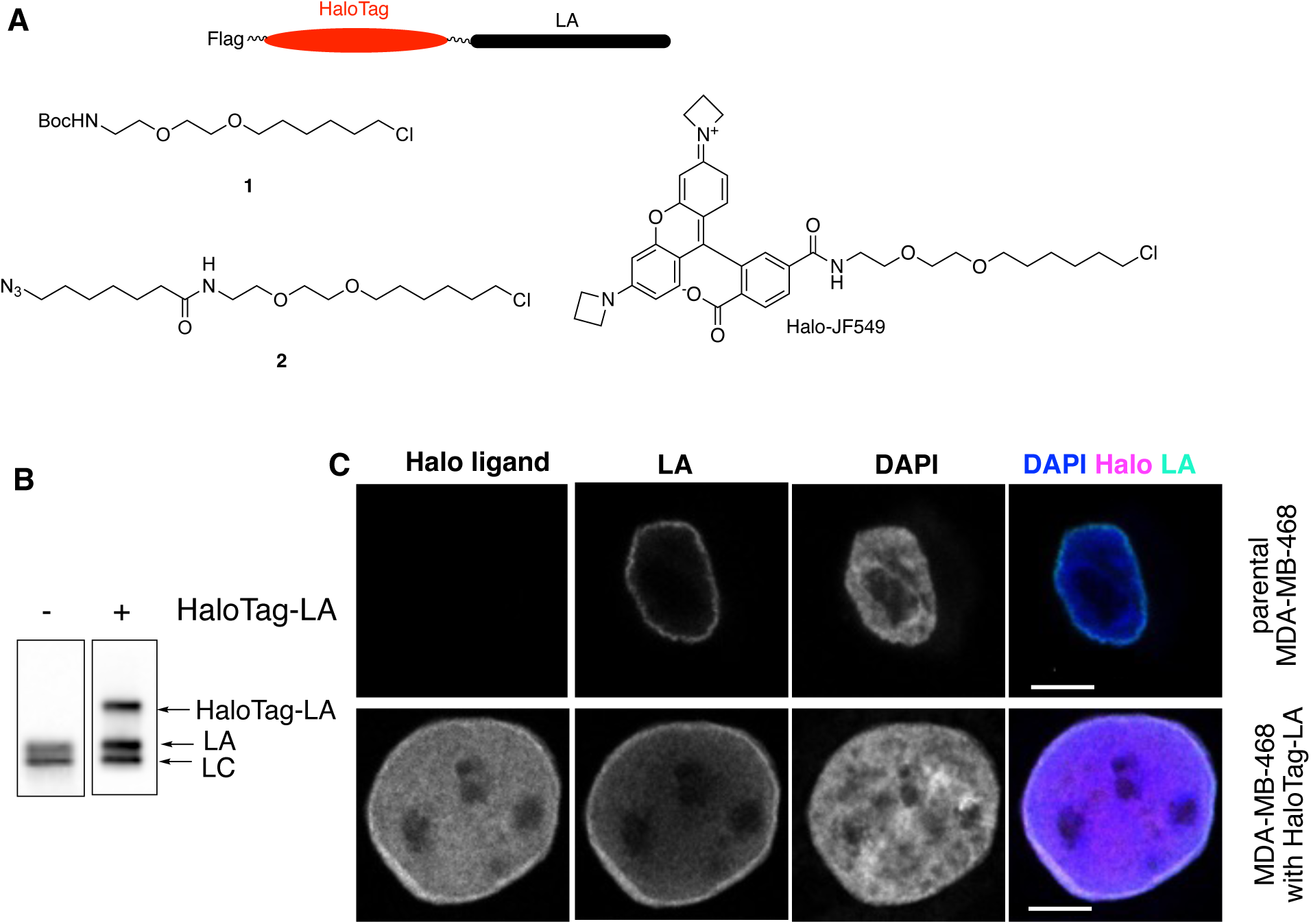
HaloTag-LA was stably expressed in MDA-MB-468 cells. (A) Schematic drawing of HaloTag-LA fusion (see Figure S1 for sequence). Chemical structures of masking halo ligand **1**, clickable halo ligand **2** and Halo-JF549 are shown. (B) HaloTag-LA was expressed in MDA-MB-468 cells. The crude cell lysates from parental cells and the cells with lentiviral expression were prepared for Western blot analysis and blotted with anti-LA. (C) HaloTag-LA fusion was stably integrated into nuclear lamina. The parental MDA-MB-468 cells and their counterparts expressing HaloTag-LA were treated with Halo-JF549. The cells were also co-stained with anti-LA. The nucleus was stained with DAPI. The confocal micrographs of representative cells are shown. Scale bars are 5 μm.

**Figure 2.**
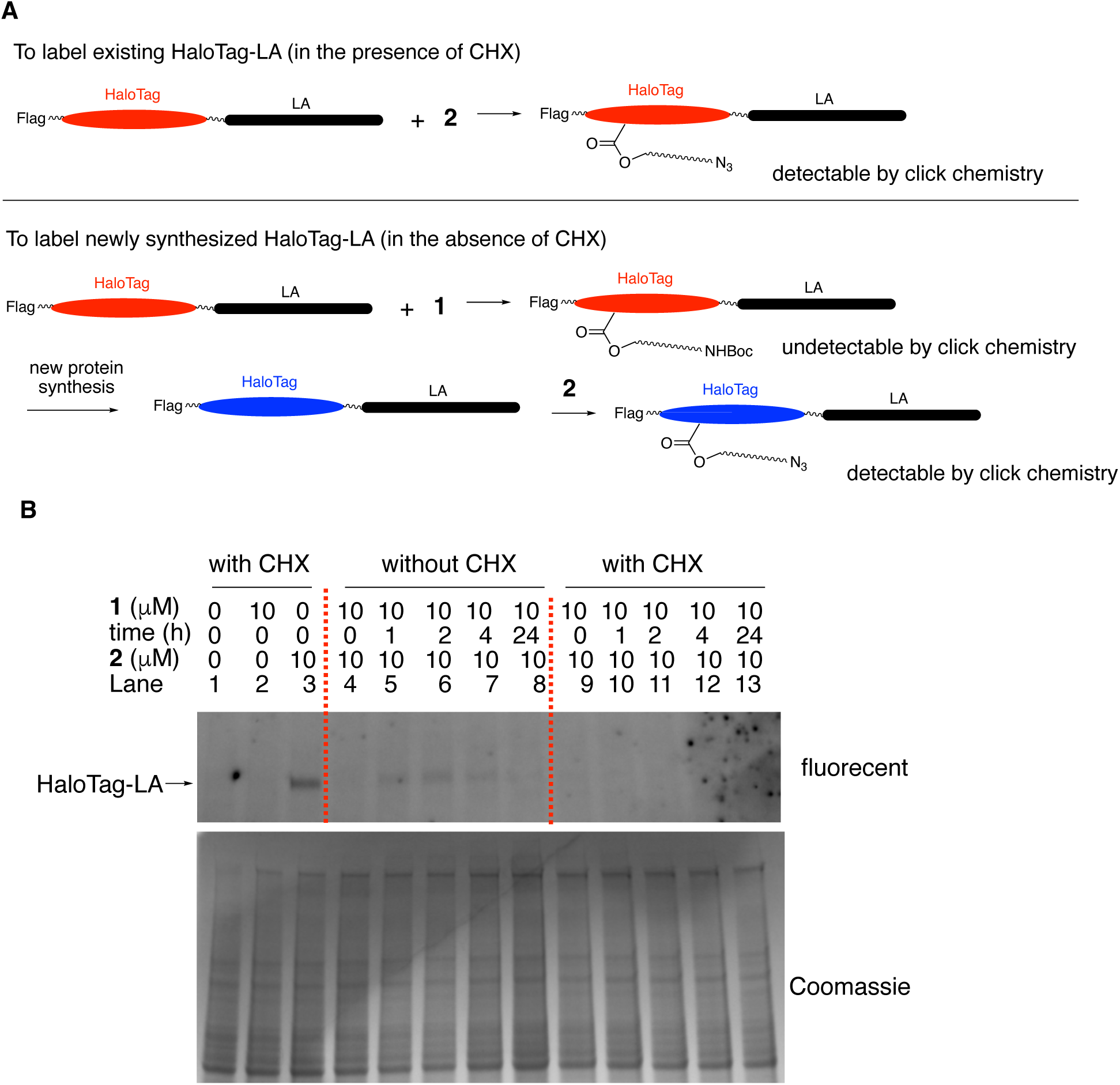
_CG_-SLENP to distinguish existing HaloTag-LA and newly synthesized HaloTag-LA. (A) Schematic diagrams to illustrate the _CG_-SLENP strategy to label existing HaloTag-LA (top) and newly synthesized HaloTag-LA (bottom). (B) MDA-MB-468 cells expressing HaloTag-LA were treated with CHX along with compound **1** for 30 min. Then the media were removed and fresh media containing CHX was added for the indicated time periods, when compound **2** (10 μM) was added and incubated for 30 min. The cells were harvested and the lysates were prepared for click reaction with TAMRA-alkyne. The resulting lysates were separated on a SDS-PAGE gel and the gel was fluorescently imaged (top). The gel was then stained with Coomassie blue (bottom) as a loading control.

With this unique HaloTag-LA system established, we interrogated the possibility to selectively detect existing HaloTag-LA and newly synthesized HaloTag-LA. To distinguish these two populations, two different Halo ligands **1** and **2** were designed (Figures 1A and 2A). Ligand **1** was designed as a masking Halo ligand while **2** was designed as a reporting ligand by utilizing the built-in bioorthogonally reactive N_3_ group, which can be detected by Cu(I)-catalyzed azide alkyne cycloaddition reaction (aka click reaction) with a tagged alkyne.^21^ The synthesis of these ligands is shown in **Scheme 1** and was adapted from a previously reported synthesis.^22^ Briefly, the Boc-protected alcohol **3** was alkylated with 1-chloro-6-iodohexane (**4**) in the presence of NaH to give Halo ligand **1**. The Boc group in **1** was deprotected under acidic condition (HCl.E_2_O) to generate amine **5**, which was further coupled with azido acid **7** to be prepared from bromo acid **6** to furnish desired Halo ligand **2**.

**Scheme 1.**
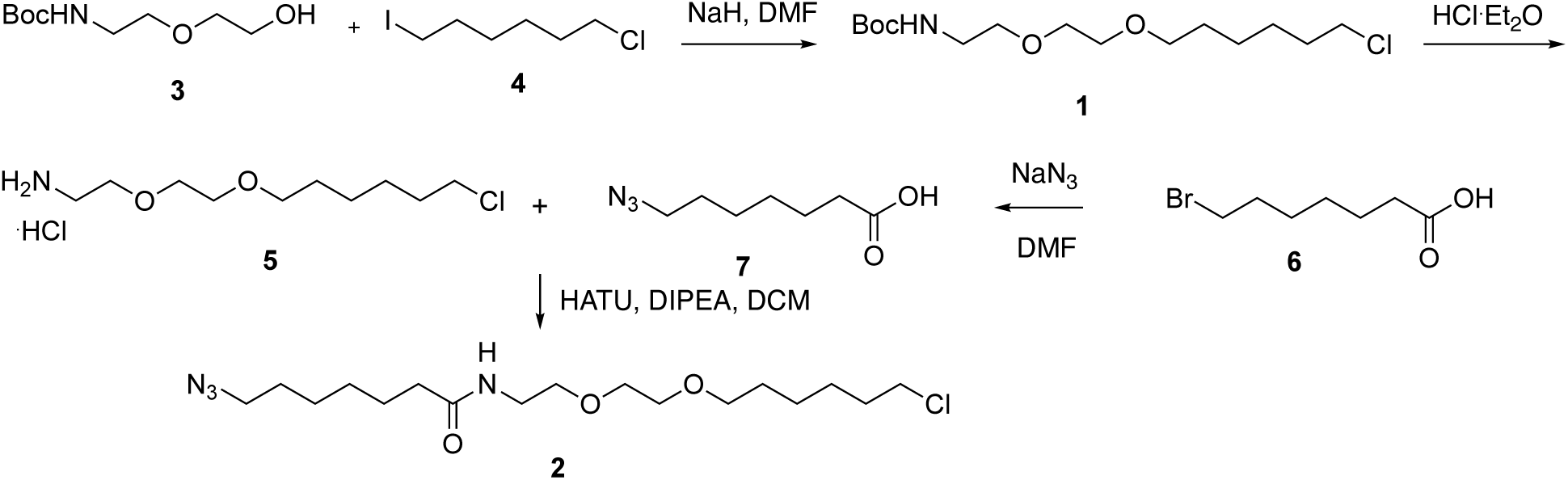
Synthesis of Halo ligands 1 and 2.

With these two different Halo ligands in hand, we envisioned that existing HaloTag-LA would be labeled by Halo ligand **2** in the presence of a protein synthesis inhibitor cycloheximide (CHX)^23^ to block new protein synthesis (Figure 2A). On the other hand, newly synthesized HaloTag-LA would be selectively labeled by first masking the existing HaloTag-LA using silent Halo ligand **1** followed by chasing the newly synthesized HaloTag-LA using Halo ligand **2**. Once the newly synthesized HaloTag-LA is labeled by **2**, it can be visualized by a click reaction with a fluorescent alkyne (Figure 2A). To test this strategy, MDA-MB-468 cells expressing HaloTag-LA were treated with CHX, then the cells were labeled with **2** (10 μM) for 30 min. The cells were harvested and the lysates were prepared for a click reaction with a tetramethylrhodamine (TAMRA) conjugated terminal alkyne in the presence of TCEP, TBTA and CuSO_4_.^24^ Then the lysates were separated on an SDS-PAGE and the gel was directly visualized by in-gel fluorescence scanning. As shown in Figure 2B, treatment with **2** resulted in specific and prominent labeling of HaloTag-LA (lane 3). On the other hand, treatment of the cells with the masking ligand **1** (10 μM) did not produce any signal (lane 2).

Pretreatment of the cells with **1** followed by labeling with **2** in the presence of CHX also produced no labeling of HaloTag-LA (lanes 9-13), indicating complete saturation of existing Halo-binding sites by **1** and no new HaloTag-LA was being synthesized under these conditions. When CHX was omitted from the treatment, we observed gradually increased labeling of HaloTag-LA when the cells were pretreated with **1** followed by a chasing period of **2** (lanes 4-7). Prominent labeling of HaloTag-LA was observed even 1 h after the existing HaloTag-LA was masked by ligand **1**, suggesting that HaloTag-LA is constantly being made in the cells. Interestingly, we observed a decrease of labeled HaloTag-LA 24 h post labeling by **2** (lane 8). The mechanism behind this decrease is presently unknown and detailed investigation was not conducted. Because HaloTag-LA is integrated into the nuclear lamina as filaments, it is possible that a fraction of Halo ligand binding site was not accessible when the cells were treated with **1**. During the subsequent period of time of labeling by **2**, the HaloTag-LA filaments might be dynamically rearranged to expose unoccupied Halo ligand binding site, which would phenotypically produce the same results even in the absence of new protein synthesis. In this case, this alternative seems unlikely because when the cells were co-treated with CHX, no additional labeling of HaloTag-LA was observed during the subsequent treatment period of time (Figure 2B, lanes 9-13). Thus, the labeling seen with the sequential treatment of **1** and **2** was indeed reporting newly synthesized HaloTag-LA. These results also suggest that the existing HaloTag-LA filaments may not undergo dynamic structural remodeling or the Halo-binding site was fully exposed for labeling by ligand **1** during the masking period. The former case is supported by previous fluorescence recovery after photobleaching (FRAP) studies with GFP-LA in different cell types to indicate that LA in the interphase is relatively immobile,^25^ which is also consistent with our live cell imaging results (*vide infra*).

With the system to differentiate existing HaloTag-LA from newly synthesized HaloTag-LA established, we employed our previously reported **LBL1** to investigate if it has differential effect on existing LA over newly synthesized LA, a property not possible to address using previously known methods. The small molecule ligand **LBL1** (Figure 3A) selectively binds to nuclear lamins in living cells.^15, 16^ Furthermore, we found that **LBL1** binds to LA(1-387) encompassing the non-helical head domain and the entire helical coiled-coil rod domain.^15^ As a coiled-coil, LA(1-387) can exist in multiple different oligomeric states in solution. As a result, in a thermal denaturing assay, we found that LA(1-387) was melting at two distinct temperatures (*T_m_*) with one melting at ∼40 °C (*T_m_*1) while the other one at ∼65 °C (*T_m_*1).^15^ Interestingly, **LBL1** only shifted *T_m_*1 without significantly affecting *T_m_*2.^15^ Based on this unique difference, we hypothesized that **LBL1** might induce formation of more oligomeric species. To test this hypothesis, we purified His-tagged LA(1-387) and investigated its oligomeric properties in the presence of **LBL1** using dimethylsuberimidate (DMS) as an amine-reactive crosslinker. DMS will react with two spatially close lysine residues.^26^ In the absence of **LBL1**, the purified LA(1-387) existed as a mixture of monomer and other oligomers as evidenced from the DMS crosslinking results (Figure 3B). When LA(1-387) was incubated with **LBL1**, more dimers, tetramers and other higher molecular weight oligomeric species of LA(1-387) were observed. These effects were dose-dependent. Interestingly, a new species at ∼75 kD was observed. This MW is slightly less than an anticipated dimer. We interpret this as a dimer with an alternative crosslinking pattern that caused its faster migration on the SDS-PAGE gel. These results support that **LBL1** can induce formation of higher molecular weight LA assemblies. They are consistent with our earlier finding that **LBL1** only stabilized the *T_m_*1 in the thermal shift assay and *T_m_*1 likely reflects the melting of oligomers whereas *T_m_*2 indicates the melting of monomer of LA(1-387).^15^

**Figure 3.**
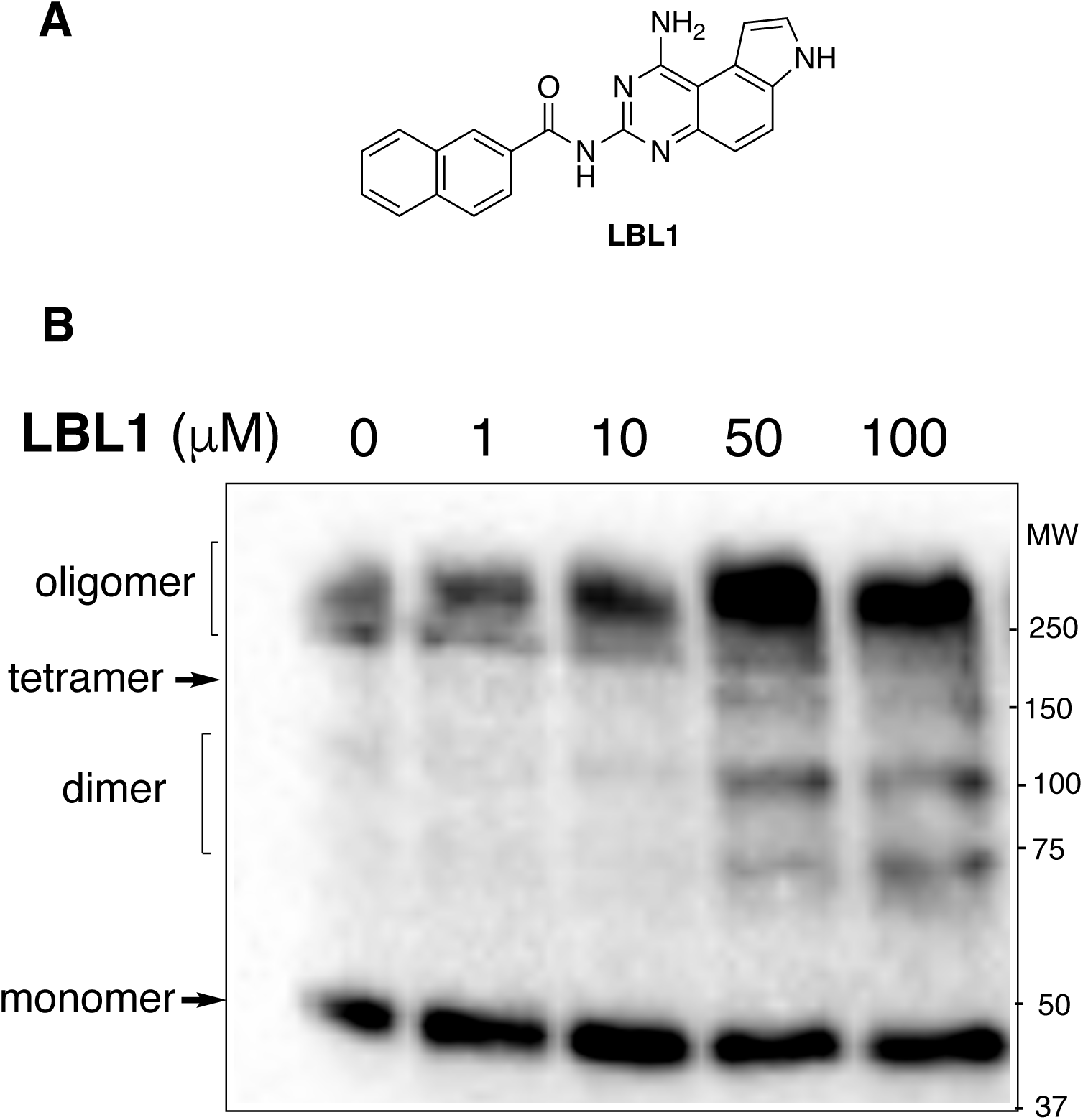
LBL1 induced formation of oligomeric LA(1-387). (A) Chemical structures of **LBL1**. (B) Recombinant His_6_-LA(1-387) was expressed and purified from *E. coli*. The purified protein was incubated with different concentrations of **LBL1**. Then DMS was added. The sample was separated on an SDS-PAGE followed by Western blotting with anti-His antibody.

The unique activity of **LBL1** towards LA(1-387) *in vitro* prompted us to investigate how **LBL1** might affect the oligomeric assembly states of existing LA and newly synthesized LA in cells using _CG_-SLENP. When HaloTag-LA molecules are assembled as filaments, it is anticipated that some of the Halo ligand binding sites might be inaccessible to Halo ligands. If **LBL1** could affect the dynamics of HaloTag-LA filament, additional binding sites will be exposed to be labeled by Halo ligand **2**. To test the susceptibility of newly synthesized HaloTag-LA protein, we first treated the cells with Halo ligand **1** to block all the available Halo binding sites in existing HaloTag-LA protein. Then the cells were treated with **LBL1** at different concentrations (0, 5, 10 μM) for different periods of time. After **LBL1** treatment, the cells were further treated with Halo ligand **2** to label any newly synthesized or exposed Halo-binding sites. These new Halo ligand binding sites could be detected by a click reaction with TAMRA-alkyne. As shown in Figure 4A and consistent with Figure 2B, the newly synthesized HaloTag-LA during the 1 h treatment period was readily detected (lane 1) by fluorescent gel scanning. Importantly, we observed increased Halo labeling upon treatment with **LBL1** (5 or 10 μM, lanes 2 and 3). The increased labeling of HaloTag-LA by Halo ligand **2** during the **LBL1** treatment period was not due to increased synthesis of HaloTag-LA caused by **LBL1** because the total HaloTag-LA amount was not changed by **LBL1** treatment as indicated by Western blot analysis using anti-Flag antibody. Similar increase of HaloTag-LA labeling by Halo ligand **2** was also observed during 2 h treatment with **LBL1** (lane 4-6, Figure 4A). These results suggest that **LBL1** could modulate the accessibility of Halo binding site in the assembled HaloTag-LA filaments. They aslo suggest that the newly synthesized HaloTag-LA is present in the cells as different filamentous structures that are uniquely sensitive to **LBL1**. To investigate **LBL1**’s effect on existing HaloTag-LA in the cells, we treated the cells with CHX to block new protein synthesis. At the same time, the cells were also treated with different concentration of **LBL1** for 1 or 2 h. Then the cells were labeled with Halo ligand **2**. The results are shown in Figure 4B. In contrast to the newly synthesized HaloTag-LA, no apparent increase of HaloTag-LA labeling by Halo ligand **2** was observed with increasing concentrations of **LBL1** either after 1 or 2 h treatment. Further prolonging the treatment to 24 h did not result in increased HaloTag-LA labeling by **2** (Figure S2). Together with the results shown in Figure 2B, these data support that the existing HaloTag-LA is well-integrated into the nuclear lamina and its assembly state is not readily affected by **LBL1** treatment. On the other hand, the newly synthesized HaloTag-LA is very different from existing ones in that their structural assembly is more readily affected by **LBL1** resulting in more Halo-binding sites being exposed for Halo ligand labeling.

**Figure 4.**
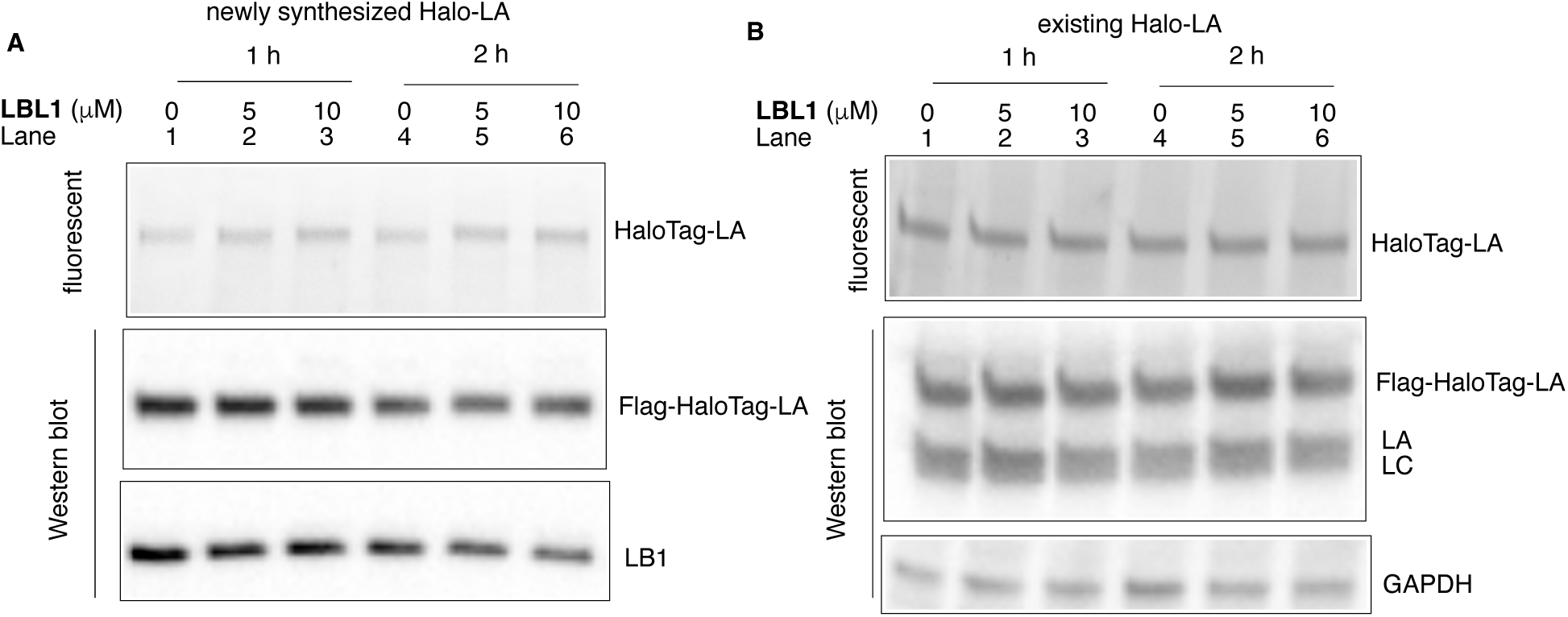
LBL1 differentially modulates the dynamics of existing HaloTag-LA and newly synthesized HaloTag-LA. (A) **LBL1** increased Halo binding sites in newly synthesized HaloTag-LA. The cells were treated with Halo ligand **1** (10 μM) for 30 min, when the media were removed and the cells were treated with **LBL1** for 1 or 2 h at the indicated concentrations. The cells were further treated with Halo ligand **2** (10 μM) for 30 min. The lysates were prepared for click reaction with TAMRA-alkyne and separated on SDS-PAGE for both fluorescent gel scanning and Western blot analyses with indicated antibodies. LB1 was used as a loading control. (B) **LBL1** did not modulate the Halo binding sites of existing HaloTag-LA proteins. The cells were treated with CHX along with **LBL1** for 1 or 2 h, when Halo ligand **2** (10 μM) was added and incubated for 30 min. The resulting cell lysates were analyzed on an SDS-PAGE after being clicked with TAMRA-alkyne. GAPDH was used as a loading control.

To further investigate the differential properties of existing HaloTag-LA and newly synthesized HaloTag-LA in live cells, we resorted to live cell imaging enabled by the fluorescent Halo ligand (Figure 5A). We first investigated the dynamics of existing HaloTag-LA by treating the cells with CHX. Then the cells were labeled with Halo-JF549 for live cell imaging. In the absence of **LBL1** treatment, HaloTag-LA is localized inside the nucleus with presence in both nuclear lamina and nucleoplasm (Figure 5B and Figure S3A). The dynamics of HaloTag-LA was monitored for 3 minutes by capturing an image every 2 seconds. As expected, the HaloTag-LA signal was pretty much immobile (Figure 5B and supplementary movie S1). When the cells were treated with **LBL1**, the HaloTag-LA was also predominantly present in the nucleus (Figure 5C and Figure S3B), which is similar to the one observed without **LBL1** treatment. However, a minor population of HaloTag-LA was detected in the cytosol (red circles in Figure 5C). While the HaloTag-LA in the nucleus was essentially immobile, the minor population in the cytosol was very dynamic and moved rapidly (red circles in Figure 5C and supplemental movie S2).

**Figure 5.**
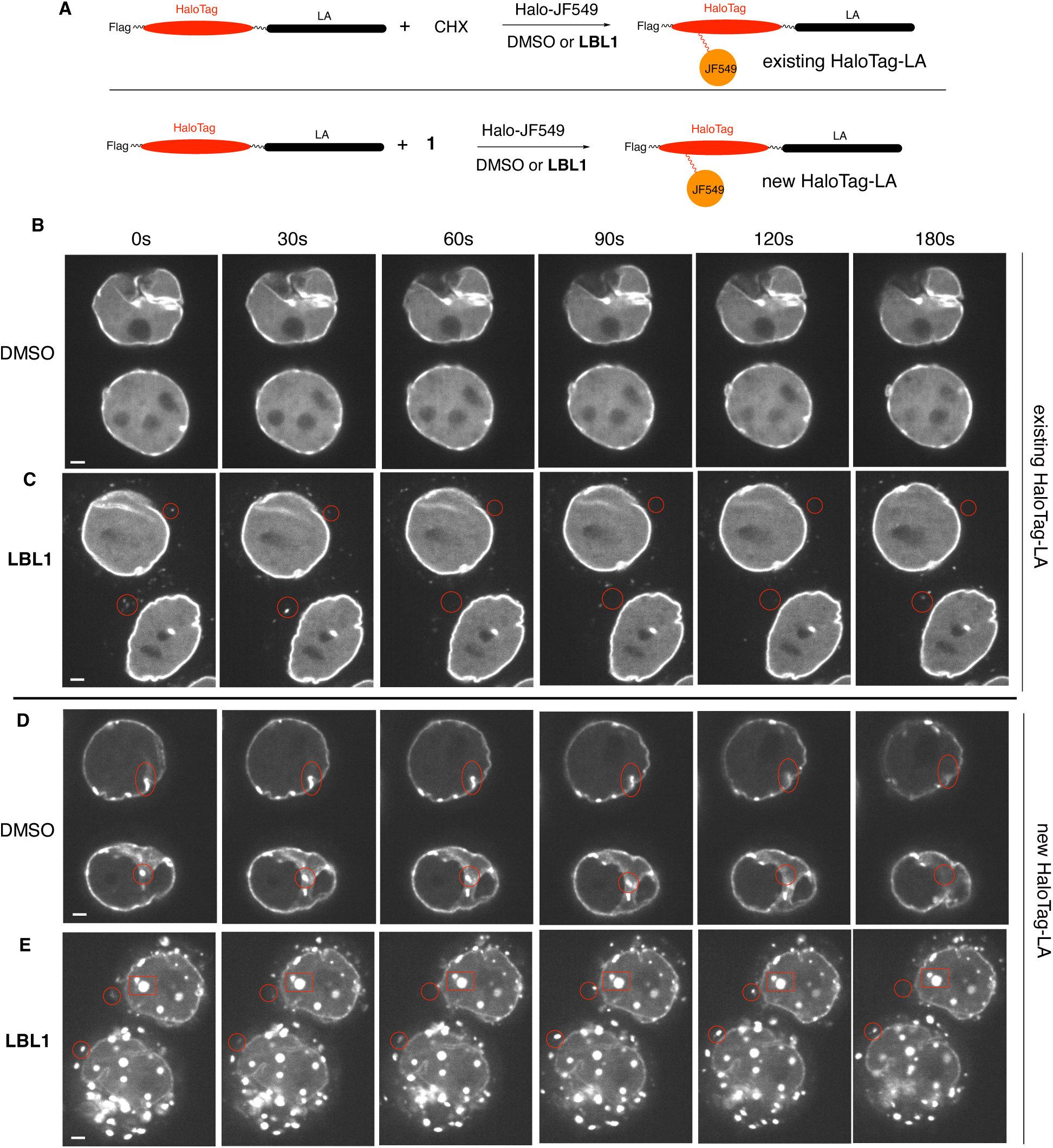
Live cell imaging of existing HaloTag-LA and newly synthesized HaloTag-LA using _CG_-SLENP. (A) Labeling schemes to selectively label existing and newly synthesized HaloTag-LA for live cell imaging. (B-C) Live cell imaging of existing HaloTag-LA. MDA-MB-468 cells stably expressing HaloTag-LA were treated with CHX in the absence (B) or presence (C) of **LBL1**. Then HaloTag-LA was labeled with Halo-JF549. The cells were then subjected to imaging and monitored for 3 minutes with an image taken every 2 seconds. (D-E) Live cell imaging of newly synthesized HaloTag-LA. MDA-MB-468 cells stably expressing HaloTag-LA were treated with compound **1** to mask existing HaloTag-LA. Then the cells were treated with DMSO (D) or **LBL1** (E). Then the newly synthesized HaloTag-LA was labeled with Halo-JF549. The cells were then subjected to imaging and monitored for 3 minutes with an image taken every 2 seconds. The fluorescence display scales for each panel were adjusted differently for better clarity. The time lapse movies corresponding to each panel are provided as online supplementary materials. The scale bars are 2 μm.

Using the masking Halo ligand **1**, we also investigated the localization and dynamics of newly synthesized HaloTag-LA in live cells (Figure 5A, bottom). We first treated the cells with Halo ligand **1** in the absence of CHX. Then the cells were allowed for new protein synthesis and assembly for 1 h. The newly synthesized HaloTag-LA was then labelled with Halo-JF549 for live cell imaging. As shown in Figure 5D, Figure S3C and consistent with Figure 2B, the newly synthesized HaloTag-LA was readily detectable, albeit at lower fluorescence intensity compared to existing ones. The newly synthesized HaloTag-LA was exclusively localized in the nucleus with prominent nuclear lamina signal, suggesting that synthesis and nuclear translocation of HaloTag-LA occurred rapidly during the 1 h period of time. In contrast to the existing HaloTag-LA, the newly synthesized HaloTag-LA displayed a few distinct characteristics. First, the newly synthesized HaloTag-LA was primarily localized at the nuclear lamina. Although nucleoplasmic HaloTag-LA was present, it was much less than what was observed with existing HaloTag-LA. Second, the localization pattern for HaloTag-LA in the nuclear lamina is less homogenous than the existing HaloTag-LA. Third, there was apparently more aggregated HaloTag-LA, which are presented as bright dots (Figure 5D and Figure S3C). Fourth, the newly synthesized HaloTag-LA was more mobile and dynamic than existing HaloTag-LA (red circle and oval in Figure 5D and also supplementary Movie S3). These features suggest that the newly synthesized HaloTag-LA has different assembly states when compared to existing HaloTag-LA in live cells.

Given these potential differences and the biochemical effect of **LBL1** on the purified LA(1-387) (Figure 3), we anticipated that **LBL1** could have different effect on the newly synthesized HaloTag-LA in live cells. To investigate this point, we treated the cells with **LBL1** after the existing HaloTag-LA was masked by Halo ligand **1**. Then the newly synthesized HaloTag-LA was labelled with Halo-JF549 for live cell imaging. As presented in Figure 5E, Figure S3D and supplementary Movie S4 and in stark contrast to existing HaloTag-LA, the localization pattern of newly synthesized HaloTag-LA was dramatically affected by **LBL1**. **LBL1** induced the formation of numerous HaloTag-LA aggregates both inside the nucleus and the cytosol. While the aggregates inside nucleus were relatively immobile (red rectangle in Figure 5E), the ones in the cytosol were very mobile and dynamic (red circles in Figure 5E). Despite these changes to the aggregates in the cytosol and nucleoplasm, **LBL1** did not induce much change to the distribution of HaloTag-LA in the nuclear lamina and this pool of HaloTag-LA was also relatively immobile. We interpret the presence of cytosolic aggregates as the effect of **LBL1** being able to inhibit the nuclear translocation of lamin protein. Based on this finding, the small cytosolic aggregates present in Figure 5C might also be due to the effect of **LBL1** to block the nuclear entry of residual lamin protein molecules that were yet to reach the nucleus.

## CONCLUSIONS

In conclusion, we developed the first chemical genetics-based SLENP approach to selectively label newly synthesized and existing proteins. This approach is potentially generalizable to any protein of interest by creating different HaloTag in-frame fusions and the application of differentially conjugated Halo ligands. While our results are mainly based on clickable azide Halo-ligand and fluorescent Halo ligands, it is anticipated that the combination of CHX and Halo-ligands with different fluorophores or clickable groups will allow multiplex labeling of newly synthesized proteins and existing ones. Using LA as an example of filament proteins undergoing multi-subunit assembly and **LBL1** as a unique small molecule to target LA, we further provide data to show that newly synthesized LA has differential susceptibility, in comparison to existing one, to small molecule ligand **LBL1**. Given the high specificity of **LBL1** for lamin labeling in cells,^16^ our results indicate that the existing LA filaments do not undergo dynamic remodeling while the new ones are more susceptible to undergo dynamic remodeling and/or subunit exchange. Consistent with our *in vitro* crosslinking results, the newly synthesized LA proteins are likely present in a lower-order assembly state and **LBL1** can selectively induce this pool to form higher-order aggregates. These distinct aggregates are reminiscent of the aggregates formed by certain LA mutants when expressed in cells.^12, 27^ These results further indicate that newly synthesized LA is at least partially structured for **LBL1** binding. This is consistent with the prevailing view of folding of nascent proteins at the ribosomes.^28, 29^ Based on the differential effect of **LBL1**, our results indicate that newly synthesized LA is structurally different from existing LA. While the details of the differences are unknown at present, the likelihood of co-translational assembly LA polypeptides and their association with mRNA during nascent protein synthesis can potentially provide unique structural features that can be differentially modulated by **LBL1**. The _CG_-SLENP approach described here provides a novel platform to selectively investigate the properties of newly synthesized proteins and existing ones.

### Experimental Section

**Chemistry and chemicals**. The details of the synthesis of Halo ligand **1** and **2** are presented in the Supplementary Materials. **LBL1** was prepared according to our previously published procedure.^30^ TAMRA-alkyne was obtained from Click Chemistry Tools (Scottsdale, AZ). Halo-JH549 were obtained from Promega (Madison, WI).

**Cell lines and culture**. HEK293T cells were purchased from ATCC. MDA-MB-468 cells were obtained from National Cancer Institute Development Therapeutic Program. The cells were authenticated by STR profiling and tested for mycoplasm contamination by PCR. The cells were routinely cultured in high glucose Dulbecco’s modified Eagle’s medium (DMEM, ThermoFisher) supplemented with 10% FBS (Hyclone) and 10% nonessential amino acids (ThermoFisher) at 37 °C with 5% CO_2_. The cells were used within 50 passages.

**Plasmids and antibodies**. Flag-tagged HaloTag-LA was designed to contain N-terminal Flag tag, HaloTag and LA. The designed in-frame fusion was assembled through gene synthesis at Genewiz (South Plainfield, NJ) and cloned into 3^rd^-generation lentiviral vector pLJM1 (Addgene). The following antibodies are used: rabbit anti-LB1 (Cell Signaling Technology), mouse M2, mouse anti-LA (Sigma), mouse anti-GAPDH (Santa Cruz Biotechnology) and mouse anti-His tag (AnaSpec). Recombinant LA(1-387) was purified as described previously.^15^

***In vitro* LA(1-387) protein crosslinking**. His-LA(1-387) in HBS (20 mM HEPES, 250 mM NaCl, pH 8.0) (40 μg/mL) was incubated with different concentrations of **LBL1** for 1 h at room temperature. Then dimethyl suberimidate (DMS, 11 mM) was added and the mixture was incubated at room temperature for 15 min, when the crosslinking reaction was quenched with addition of glycine (50 mM). The mixture was incubated at room temperature for 10 min. SDS-PAGE buffer was added and the sample was subjected to Western blot analysis using anti-His antibody.

**Lentivirus preparation and transduction**. The lentivirus was prepared as previously described.^31^ Briefly, lentiviruses expressing HaloTag-LA were prepared from HEK 293T cells by co-transfecting lentiviral expression plasmid along with packaging vectors using the calcium-phosphate method (TaKaRa). MDA-MB-468 cells were transduced with the prepared lentiviruses expressing HaloTag-LA. After puromycin selection, the cells were used for subsequent experiments.

**Confocal microscopy**. This procedure is similar to previously published method^24^ with slight modifications. MDA-MB-468 cells stably expressing HaloTag-LA and parental MDA-MB-468 cells were cultured in DMEM with 10% FBS. Once the cells reached ∼80% confluence, they were harvested and transferred to coverslips, which were coated with poly-D-Lysine (R&D systems) for 24 hr. The cells were then treated with 2 µM Halo-JF549 for 15 minutes. The labeling media were removed and the cells were washed once with fresh media. Then the cells were fixed using 4% paraformaldehyde in PBS for 15 minutes at room temperature followed by 10-minute permeabilization with 0.3% triton X-100 in PBS at room temperature. The cells were blocked in 3% BSA in PBS for one hour at room temperature followed by overnight incubation with anti-LA antibody (1:8000 in PBS with 3% BSA) at 4 °C. The next day cells were stained with a secondary Alexa fluor 488 conjugated anti-mouse antibody (Jackson ImmunoResearch Laboratories) for one hour at room temperature. The cells were further incubated with 300 nM DAPI for 10 minutes. Images were taken on an inverted Zeiss LSM 980 confocal microscope.

_CG_**-SLENP and click chemistry**. MDA-MB-468 cells stably expressing HaloTag-LA were treated with or without CHX (0.1 mg/mL) along with Halo ligand **1** (10 μM) for 30 min. The media were removed and the cells were washed once with media. Then the cells were treated with indicated concentrations of **LBL1** for different periods of time. Then Halo ligand **2** (10 μM) was added for another 30 min of labeling. The cells were harvested and washed twice with ice-cold PBS. The cell pellets were lysed in 1%SDS in PBS with sonication. Then equal amount of protein lysates (∼10 μg) was clicked with TAMRA-alkyne (10 μM) and TBTA (100 μM), TCEP (1 mM), and CuSO_4_ (1 mM) for 1.5 h at room temperature.^32^ The final SDS concentration in the click reaction was ≤ 0.5%. Then the proteins were separated on a 4-20% precast SDS-PAGE gel (Biorad). The gel was scanned for fluorescent signal on ChemiDoc^MP^ (Biorad) before it was transferred to a nitrocellulose membrane (Biorad) for western blotting with indicated antibodies.

_CG_**-SLENP for live cell imaging**. MDA-MB-468 cells stably expressing HaloTag-LA were cultured with FluoroBrite^TM^ DMEM (ThermoFisher) and 10% FBS. Once cells reached ∼80% confluence, they were collected and transferred to glass bottom 4-well pie dishes (Greiner Bio-One cat# 627871), which were coated with poly-D-Lysine (R&D systems) for 24 hr. For imaging existing HaloTag-LA, the cells were treated with CHX (100 μg/ml) for one hour. Then DMSO or **LBL1** (10 μM) was added for 1 h. This was then followed by a 15-minute treatment with 2 µM of Halo-JF549 ligand (Promega). Afterward, the labeling media were removed and fresh media containing DMSO or **LBL1** (10 μM) were added. The cells were then subjected to live cell microscopic imaging. Live cell imaging was performed using an inverted Nikon TiE microscope equipped with a Yokogawa CSU-W1 spinning disk confocal head, running with NIS Elements software (Nikon). The cells were maintained at 34.5 °C and 5% CO_2_. Images were acquired with a Zyla v5.5 sCMOS camera (Andor) with a 100x (NA 1.49) objective. The Halo-JF549 was excited with a 561 nm laser and emission light was collected between 570 nm and 640 nm. A single image was taken every two seconds during each three-minute time lapse movie. For imaging newly synthesized HaloTag-LA, the cells were treated with compound **1** (10 μM) for 30 minutes, when the cells were washed six times with media and then incubated in media containing DMSO or **LBL1** (10 μM) for one hour. Next, the cells were labeled with 2 µM of Halo-JF549 for 15 minutes. The labeling media were removed and fresh media containing DMSO or **LBL1** (10 μM) were added. The cells were then subjected to live cell microscopic imaging as above.

## Acknowledgements

This work was made possible by financial supports provided from R21EB028425 (BXL), R01CA245964 (BXL), R01CA278058 (BXL and XX), R01CA211866 (XX), and R01GM122820 (XX). We thank OHSU Gene Profiling Shared Resource for authenticating the cell lines through STR profiling and OHSU Advanced Light Microscopy Shared Resource for providing technical support. The OHSU Advanced Light Microscopy Shared Resource and Gene Profiling Shared Resource were partially supported by P30CA069533.

